# The role of pneumococcal extracellular vesicles on the pathophysiology of the kidney disease Hemolytic Uremic Syndrome

**DOI:** 10.1101/2023.01.30.526387

**Authors:** Miriana Battista, Bianca Hoffmann, Yann Bachelot, Lioba Zimmermann, Laura Teuber, Aurélie Jost, Susanne Linde, Martin Westermann, Mario M. Müller, Hortense Slevogt, Sven Hammerschmidt, Marc Thilo Figge, Cláudia Vilhena, Peter F. Zipfel

## Abstract

*Streptococcus pneumoniae*-induced hemolytic uremic syndrome (Sp-HUS) is a kidney disease characterized by microangiopathic hemolytic anemia, thrombocytopenia, and acute kidney injury. This disease is frequently underdiagnosed and its pathophysiology is poorly understood. In this work, we compared clinical strains, isolated from infant Sp-HUS patients, to a reference pathogenic strain D39, for host cytotoxicity and further explored the role of Sp-derived extracellular vesicles (EVs) in the pathogenesis of a HUS infection. In comparison with the WT strain, pneumococcal HUS strains caused significant lysis of human erythrocytes and increased the release of hydrogen peroxide. Isolated Sp-HUS EVs were characterized by performing dynamic light-scattering microscopy and proteomic analysis. Sp-HUS strain released EVs at a constant concentration during growth, yet the size of the EVs varied and several subpopulations emerged at later time points. The cargo of the Sp-HUS EVs included several virulence factors at high abundance, i.e., the ribosomal subunit assembly factor BipA, the Pneumococcal Surface Protein A (PspA), the lytic enzyme LytC, several sugar utilization and fatty acid synthesis proteins. Sp-HUS EVs strongly downregulated the expression of the endothelial surface marker PECAM-1 and were internalized by human endothelial cells. Sp-HUS EVs elicited the release of pro-inflammatory cytokines (IL-1ß, IL-6) and chemokines (CCL2, CCL3, CXCL1) by human monocytes. These findings shed new light on the overall function of Sp-EVs, in the scope of infection-mediated HUS, and suggest new avenues of research for exploring the usefulness of Sp-EVs as therapeutic and diagnostic targets.

**Importance:** *Streptococcus pneumoniae* is a life-threatening human pathogen associated with severe illnesses in the upper respiratory tract. Disseminated infections also occur, as the kidney disease hemolytic uremic syndrome. Even though vaccination is available, this pathogen is responsible for a worldwide high mortality rate, especially among children from least developed countries, where vaccination strategies are poor or inexistent. It is estimated that 30% of invasive pneumococcal diseases are caused by antibiotic resistant bacteria, leading to the classification of “serious threat” by the World Health Organization. In order to prevent cases of severe illness, investigation in the direction of new vaccine candidates is of upmost importance. Pneumococcal extracellular vesicles pose as ideal candidates for a serotype-independent vaccine formulation. To this purpose, the aspects of vesicle formation, cargo allocation and function need to be understood in detail.

**Graphical Abstract:** 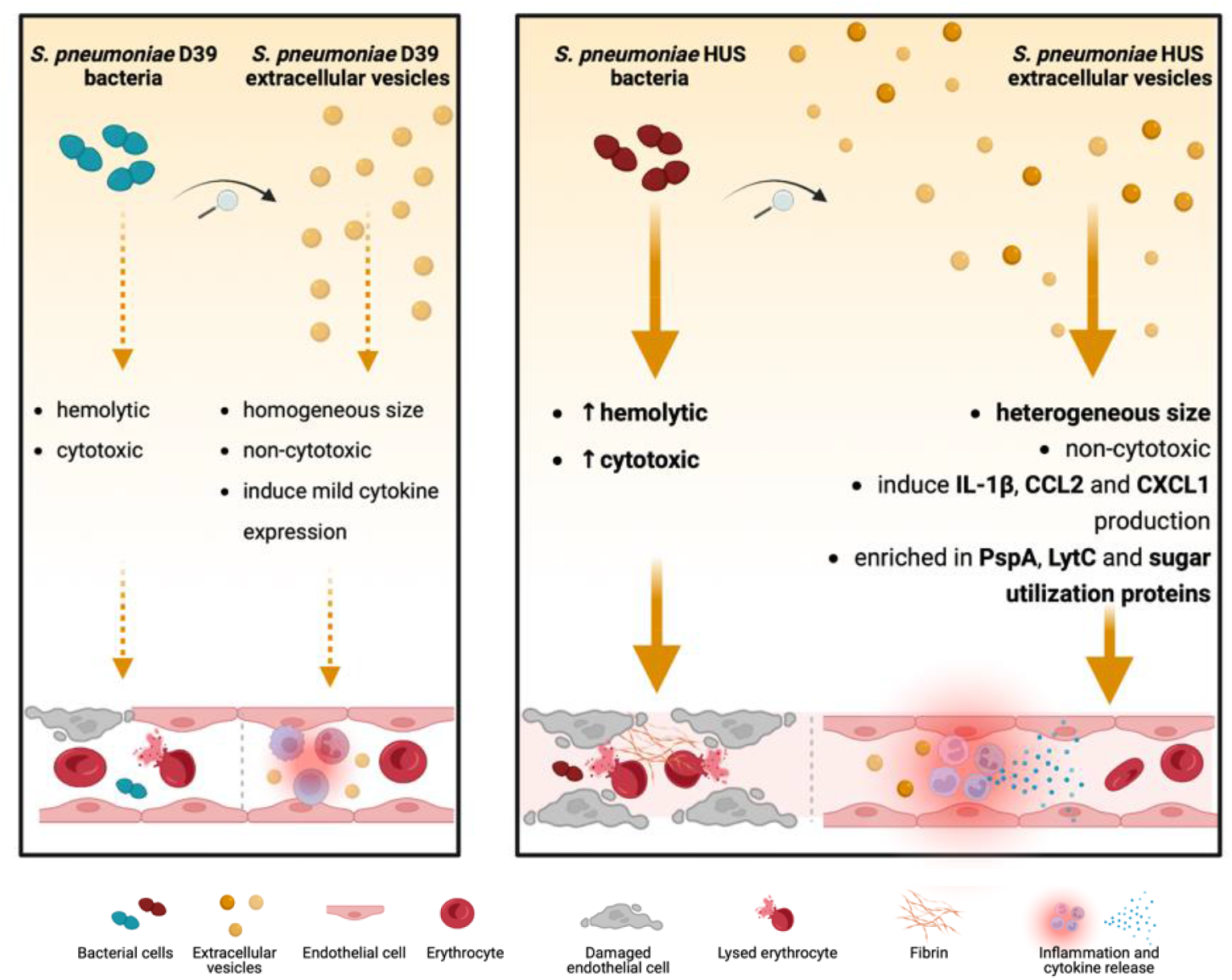

## Introduction

Hemolytic uremic syndrome caused by *Streptococcus pneumoniae* (Sp-HUS) is a rare and serious infection-induced kidney disease, clinically defined by microangiopathic hemolytic anemia, thrombocytopenia, acute kidney failure(1), and endothelial injury(2). The pathophysiology of genetic and autoimmune forms of HUS are relatively well understood. Defective complement action results in endothelial damage, thrombus formation, and, ultimately, in occlusion of small vessels in the kidney(3). However, the pathophysiology of the infection-associated form of HUS remains largely unclear (4). In around 90% of cases, an infection-related HUS is caused by Shiga-like toxin-producing bacteria, such as enterohaemorrhagic *Escherichia coli* (STEC) or *Shigella dysenteriae* type 1(5). Sp-HUS accounts for approximately 5% of all HUS cases and occurs mainly among children under 2 years old, and the predicted mortality rate is 12.3%(6, 7). The prevalence of this disease and its severe outcomes, if untreated, desperately argue for investigation of its pathophysiology.

*Streptococcus pneumoniae* is a Gram-positive human pathogen capable of causing otitis media, sinusitis, community-acquired pneumonia, and serious disseminated diseases such as meningitis and septicemia, following colonization of the upper respiratory tract. Despite the implementation of pneumococcal polysaccharide and conjugated vaccines, which confer protection only against a defined number of capsular serotypes, *S. pneumoniae* is still the leading cause of mortality in children under the age of five (8)(9). In the search for a serotype-independent vaccine, pneumococcal extracellular vesicles (Sp-EVs) have recently emerged as potential candidates and studies have shown that Sp-EVs and membrane particles exhibit good immunogenicity(10–13). Pneumococcal EVs have been isolated and shown to contain proteins such as penicillin-binding protein 1B (Pbp1B), neuraminidase A (NanA), pneumococcal surface adhesin A (PsaA), pneumolysin (Ply), and the pneumococcal surface protein A (PspA)(14, 15). Sp-EVs can bind to and be internalized by human macrophages, and epithelial and dendritic cells, and have demonstrated immunomodulatory capacities *in vitro,* eliciting the production of IL-10, IL-6, and TNF-α by the host(16–18).

EVs are associated with renal diseases such as acute kidney injury, glomerular, and tubular diseases(19). In the kidney, host-derived EVs are produced by blood cells, podocytes, endothelial, and tubular epithelial cells (20–23), and are associated with Shiga toxin-producing *E. coli* HUS (STEC-HUS), another infection-related HUS form(24–26). However, the role played by pathogen-EVs, e.g., Sp-EVs, in the establishment of HUS remains elusive.

In the present work, we first compared several cytotoxic features of a pneumococcal HUS clinical isolate with those of a reference strain and, subsequently, we isolated, characterized, visualized, and assayed Sp-EVs for their immunomodulatory activity on human host innate immune cells.

## Results

### Sp-HUS strain shows cytotoxicity towards human red blood and endothelial cells

Microangiopathic hemolytic anemia is a prominent clinical manifestation of HUS(27). Thus, we investigated the hemolytic activity of eight clinical *S. pneumoniae* HUS strains isolated from infant patients. When Sp-HUS strains were compared with the pathogenic reference strain D39, referred to as the wild type (WT), seven of the eight Sp-HUS strains showed stronger lysis of human erythrocytes (**Fig. 1A**). Hydrogen peroxide (H_2_O_2_) is the main mediator of pneumococcal hemolytic activity and also contributes to lung cellular damage(28, 29). Hence, the excreted H_2_O_2_ in bacterial supernatants from the clinical pneumococcal strains was quantified (**Fig. 1B**). HUS A strain showed higher H_2_O_2_ excretion (~2-fold more than WT) than the other strains assayed. Previously, we showed that HUS A strongly binds lactoferrin(30, 31). Thus, the HUS A strain (from now on referred to as the Sp-HUS strain), which was isolated from a 2-year-old patient suffering from HUS, was chosen as the focus for the rest of this study. Growth of the WT and Sp-HUS strains in rich medium showed that although both strains reached the same maximum optical density (OD), the Sp-HUS strain exhibited a shorter lag time, entering exponential growth earlier (at around 1 h) than WT (**Fig. S1**). Moreover, the exponential phase of growth is faster for the Sp-HUS strain than the WT.

**Figure 1.**
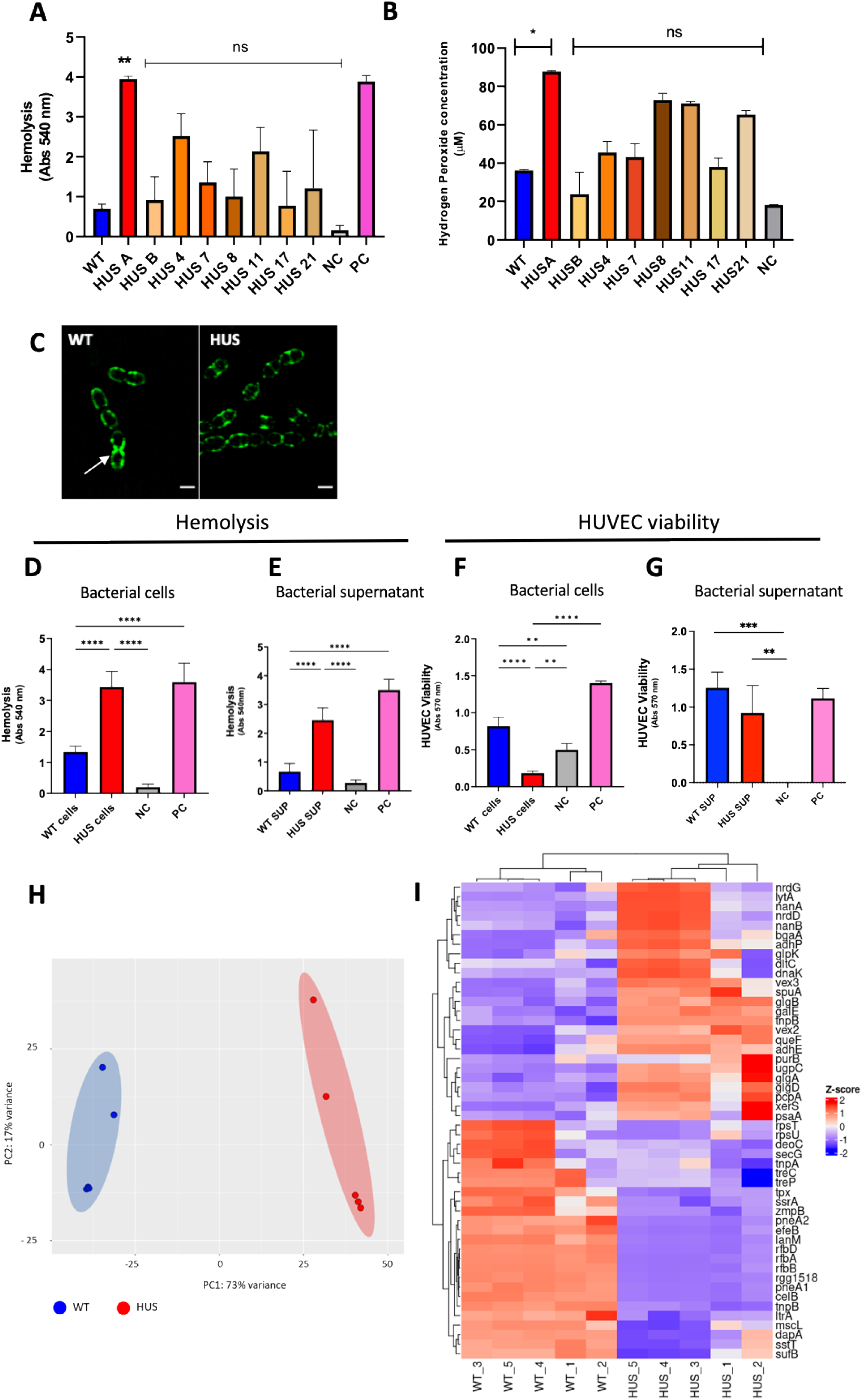
Sp-HUS strains and supernatant-mediated damage on human red blood cells and endothelia. Eight Sp-HUS strains (HUS A, B, 4, 7, 8, 11, 17 and 21) were assayed for their capacity to hemolyze human red blood cells (**A**). Red blood cells from healthy donors, were incubated with mid-exponentially grown bacterial cells at 37°C. PBS and bi-distilled water served as negative and positive controls, respectively. The absorbance of the supernatants was measured at 540 nm. Mean ± SD (**B**) Released hydrogen peroxide was quantified on the supernatant of the same strains. Bacterial cultures were harvested at mid-exponential phase, centrifuged and the supernatant immediately assayed for the presence of hydrogen peroxide using a commercial kit. (**C**) Representative SR-SIM images of WT and Sp-HUS strains stained with WGA-488. Scale bar is 1 μm. Cellular (**D**) and supernatant fractions (**E**) of selected strains were tested separately for their hemolytic capacity. (**F**) HUVECs viability was evaluated after incubation with WT or Sp-HUS strains or the corresponding supernatant fractions (**G**) by CTB assay. Tert-butyl hydroperoxide (400 μM) and DMEM were used as negative (grey bar) and positive (pink bar) controls, respectively. (**H**) PCA plot of bacterial transcriptome from RNAseq analysis. (**I**) Heatmap of differentially expressed genes from the RNAseq analysis of five replicates of WT and Sp-HUS bacteria highlighting upregulated (red) and downregulated (blue) genes in the Sp-HUS strain.

To investigate bacterial cell wall morphology, mid-exponential phase Sp-HUS and WT cultures were stained with fluorescently-labeled Wheat Germ Agglutinin (WGA), which is a carbohydrate-binding lectin with high affinity for N-acetylglucosamine (a main structural component of the pneumococcal cell-wall(32)), and stained cells were visualized using Super Resolution-Structured Illumination Microscopy (SR-SIM) (**Fig. 1C**). The distribution of WGA across the bacterial cell wall is typically homogenous, as shown on the WT panel, with exception of the division septa during later stages of cell division. At an intermediate stage of cell division, WGA dye accumulates at the septum of individual cells, as shown in the WT panel on the bottom cell (Fig. 1C arrow). The WGA-binding profile of the Sp-HUS strain differs from the WT since the Sp-HUS strain tended to grow in longer chains in which WGA frequently appeared bound to the division septum.

Next, both bacterial cells and supernatants were assessed for their hemolytic activity and influence on endothelial cell viability. The cell-associated hemolytic activity of the Sp-HUS strain was significantly higher (more than 2-fold) than the WT (**Fig. 1D**). Moreover, the supernatant fraction of the Sp-HUS strain also showed higher hemolytic activity against human erythrocytes than that of the WT (**Fig. 1E**). Since intracytoplasmic isocitrate dehydrogenase (IDH) converts resazurin into resorufin, a fluorescent end product, in intact cells, cellular metabolic activity can be monitored by appearance of this fluorescent product(33). Sp-HUS cells decreased the viability of human endothelial cells by more than 80%, consistent with their highly toxic activity in the hemolytic assay (**Fig 1F**). By contrast, however, the supernatant of neither bacterial strain decreased the viability of endothelial cells (**Fig. 1G**), which did not correlate with the hemolytic activity observed for the Sp-HUS supernatant. The effect of the Sp-HUS strain on endothelial cell retraction was also evaluated using scanning electron microscopy (SEM). An image analysis pipeline was developed using the visual programming language JIPipe(34) (steps summarized in **Fig. S2A-F**) to quantify the background fraction as an indirect measure of human cell retraction. Pneumococcal strains led to high levels of cell retraction (approx. 6-fold more than the unstimulated control). However, no significant difference was observed between Sp-HUS and the reference pathogenic strain (**Fig. S2G**).

To provide insights into relevant, cytotoxicity-related genes, the transcriptome of each strain was analyzed by RNAseq. Five biological replicates of each strain were assayed and differential gene expression (DGE) analysis was conducted using DESeq2 (35). Principal Component Analysis (PCA) validated the clusterization of replicates from each strain (**Fig. 1H**). Of the 572 genes identified, 50 genes (**Fig. 1I** and **Table 1**) matched the thresholds on expression and significance (|log_2_ FC| > 1,5 and adjusted *p*-value < 0.05). Upregulated genes in the Sp-HUS strain included genes encoding two choline-binding proteins (*lytA* and *pcpA*), two neuraminidases (*nanA* and *nanB*), genes involved in the anaerobic ribonucleotide reductase system (*nrdD* and *nrdG*), genes related to glycerol and sugar metabolism (*glpA*, *glpB*, *glpD*, *glpK*, and *galE*), and two genes involved in pyruvate metabolism to ethanol (*adhE* and *adhP*). Among the most highly downregulated genes were an iron-sulfur biogenesis system gene (*sufB*), an amino acid synthesis gene (*dapA*), a large conductance mechanosensitive channel gene (*mscL*), *ltrA*, *tnpB*, and a competence gene (*celB*). Based on the gene ontology (GO) analysis (**Fig.S3A**), the most significantly enriched GO terms referred to “organic substance metabolic/catabolic process”, “carbohydrate metabolic/catabolic process”, and “catabolic process”. Other enriched terms referred to “polysaccharide biosynthesis”, “glucan and glycogen metabolism”, and “energy derivation by oxidation of organic compounds”. A network analysis based on the GO enrichment was performed (**Fig. S3B**). Three large clusters (relating to metabolic and catabolic processes) and four minor clusters were observed. These results suggest that, in general, metabolic and catabolic related pathways are remarkably altered in the Sp-HUS strain and a complex rearrangement of carbohydrate, glycogen, and glycan metabolic pathways might contribute to the pathogenicity of this strain.

**Table 1.** Differentially expressed genes (Sp-HUS vs WT)

| Downregulated |  |  | Upregulated |  |  |
| --- | --- | --- | --- | --- | --- |
| Gene | log2FoldChange | Pvalue adj | Gene | log2FoldChange | Pvalue adj |
| rfbA | -12,18 | 2,97E-87 | nanB | 5,65 | 4,27E-8 |
| rfbB | -11,99 | 1,91E-57 | lytA | 3,14 | 2,36E-6 |
| rfbD | -11,82 | 1,83E-34 | pcpA | 3,05 | 1,75E-12 |
| rgg1518 | -9,75 | 3,00E-23 | queF | 3,00 | 2,90E-4 |
| lanM | -9,17 | 7,59E-19 | adhP | 2,82 | 7,05E-8 |
| celB | -9,16 | 7,60E-20 | nrdD | 2,80 | 1,31E-4 |
| pneA2 | -8,29 | 8,19E-14 | nrdG | 2,27 | 5,20E-3 |
| efeB | -8,18 | 2,31E-15 | galE | 2,26 | 3,64E-11 |
| pneA1 | -7,79 | 1,90E-14 | psaA | 2,09 | 6,13E-4 |
| treP | -3,77 | 2,08E-5 | nanA | 2,08 | 2,12E-5 |
| treC | -3,59 | 6,89E-6 | tnpB | 2,05 | 3,07E-19 |
| zmpB | -3,11 | 1,54E-11 | adhE | 1,94 | 3,78E-4 |
| tnpB | -2,37 | 6,61E-40 | vex3 | 1,92 | 1,25E-9 |
| ltrA | -2,30 | 1,35E-8 | glpK | 1,91 | 6,46E-3 |
| rpsT | -2,19 | 4,13E-4 | xerS | 1,90 | 3,86E-5 |
| mscL | -2,17 | 2,60E-4 | dltC | 1,89 | 1,13E-2 |
| deoC | -2,17 | 1,66E-4 | vex2 | 1,85 | 6,16E-7 |
| dapA | -1,92 | 1,41E-3 | dnaK | 1,80 | 2,07E-3 |
| secG | -1,90 | 1,93E-3 | glgA | 1,79 | 6,39E-8 |
| tpx | -1,84 | 8,88E-10 | glgD | 1,78 | 6,68E-9 |
| sstT | -1,84 | 1,02E-3 | purB | 1,72 | 2,01E-2 |
| ssrA | -1,73 | 3,28E-9 | bgaA | 1,70 | 1,29E-2 |
| rpsU | -1,70 | 8,94E-3 | ugpC | 1,51 | 2,19E-9 |
| tnpA | -1,60 | 2,72E-2 | spuA | 1,50 | 1,97E-7 |
| sufB | -1,53 | 5,06E-3 | glgB | 1,50 | 3,40E-14 |

### Heterogeneous size profile and altered protein cargo characterize Sp-HUS EVs

We showed that Sp-HUS strain causes release of hydrogen peroxide, hemolysis of erythrocytes, and reduced endothelial cell viability. Sp-HUS-derived supernatant also caused strong hemolysis, which raised the question of which supernatant components caused this effect. Pneumococcal extracellular vesicles (Sp-EVs) have important immunomodulatory capacities, affecting both human macrophages and human epithelial cells (18, 36). To address the role of Sp-HUS EVs in the interaction with host cells, Sp-EVs were isolated and characterized.

Protrusions at several subcellular locations could be observed (arrows) when *S. pneumoniae* strains were visualized using SEM (**Fig. 2A**). The protrusions or particles were visible at the bacterial septum, poles, and mid-cell, and occasionally covering the entire cell surface. These different locations relate to the bacterial cell cycle stage and the intrinsic capacity of each individual bacterium to produce these particles(15). These structures were heterogeneous in size and their round shape was suggestive of extracellular vesicles (EVs)(37–39). The cell-attached structures visualized by SEM were eventually released into the supernatant as fully-formed EVs. EVs isolated from bacterial supernatants were visualized by dynamic light-scattering microscopy, using nanoparticle tracking analysis, and the movement of the particles was followed through a microfluidic circuit. Representative snapshots of these circulating particles are shown in **Fig. 2B**. To ascertain whether the formation and size of EVs related to the bacterial growth phase, Sp-EVs were characterized over time. Histograms representative of the particle profile are depicted in **Fig. 2C** and a sum-up graph of size and concentration can be seen in **Fig. 2D** (WT-EVs) and **Fig. 2E** (Sp-HUS-EVs). WT-EVs maintain a stable size range (~120 nm) and a rather homogeneous population (unimodal histogram). The concentration increases slightly after 4 h of growth. On the other hand, the size of Sp-HUS EVs decreased over time (from ~125 nm to ~60 nm), and the population became very heterogeneous, with several subpopulations appearing after only 2 h growth, and culminated with five subpopulations at time point 4 h. By contrast, with WT-EVs, which increased in concentration during growth, the concentration of Sp-HUS EVs was stable throughout the course of growth. This suggested a difference in the regulation of EV formation between the Sp-HUS and WT strains.

**Figure 2.**
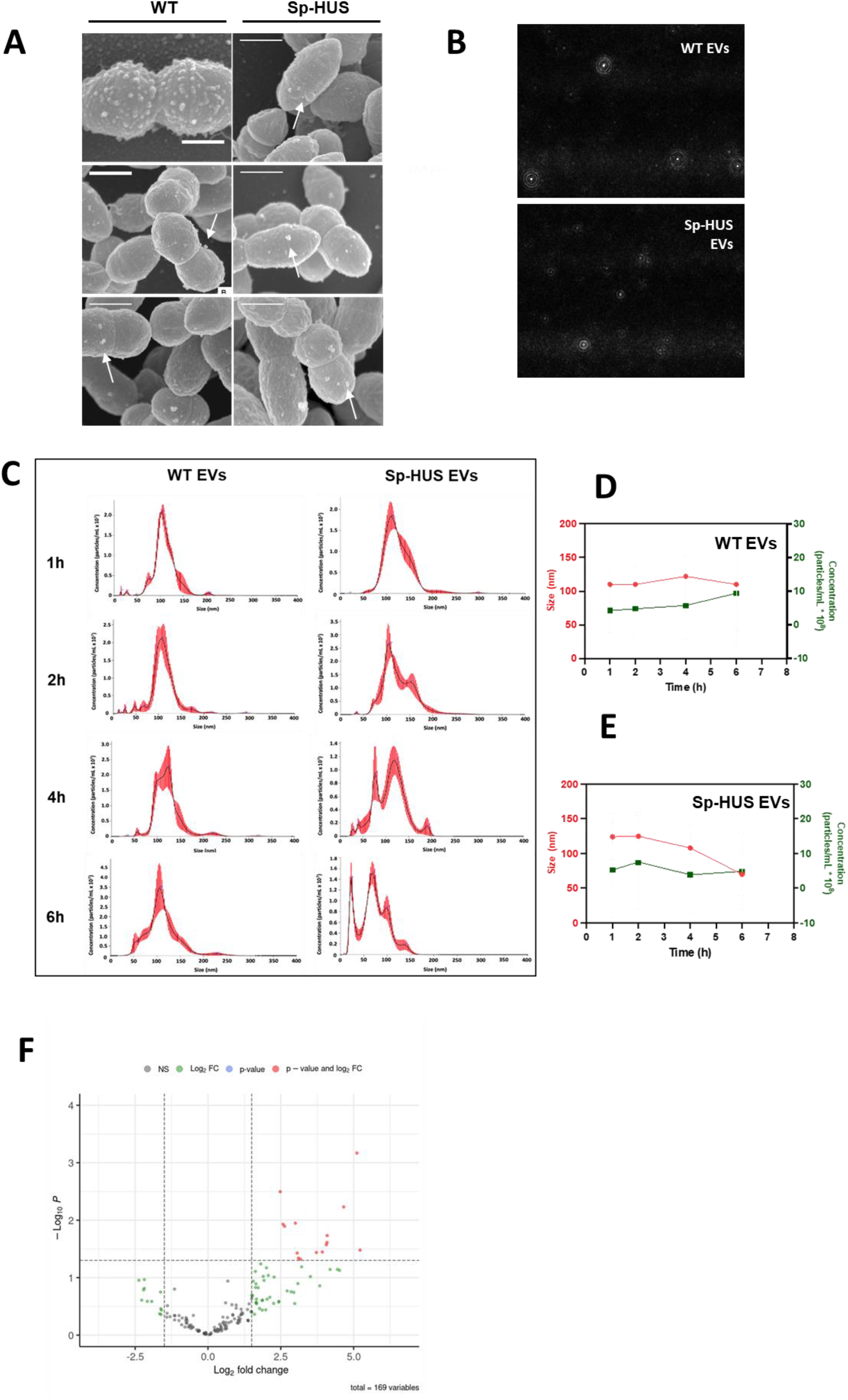
Characterization of pneumococcal EVs. **(A)** Representative SEM images of *S. pneumoniae* strains during shedding of EVs (arrows). Scale bar = 560 nm. **(B)**Representative snapshots of WT and Sp-HUS EVs, visualized by light scattering microscopy. **(C)** Histograms of EVs concentration (particles/mL x 10^7^) and size (nm), after nanoparticle tracking analysis. EVs were isolated from WT or Sp-HUS supernatant at different time points of growth (1, 2, 4 and 6 hours). **(D, E)** Sum up graphs showing WT (D) and Sp-HUS (E) EVs size (nm) (red line) and concentration (particles/mL x 10^8^) (green line) over time. **(F)** Proteomic analysis of pneumococcal EVs cargo. The proteome of Sp-HUS EVs fraction was compared to the proteome of WT EVs. Fold changes in protein levels are represented as the log_2_ ratio of Sp-HUS EVs and WT EVs (log_2_(FoldChange)), while the significance of these changes is represented by the negative logarithm of the p-value of a *t* test corrected for multiple comparisons (−log_10_ - *P* value). Significantly upregulated proteins of HUS-EVs compared to WT EVs are shown in red.

To investigate whether different production stimuli, i.e., host produced compounds, influenced vesicle concentration and size, a co-cultivation multi-well model was developed using the *Transwell* system. In this model, separate compartments, within the wells of 24-well plates, were seeded with HUVECs on the basolateral side and *S. pneumoniae* cultures on the apical side, separated by a 0.40 μm pore size membrane, allowing co-cultivation but avoiding direct contact between bacteria and host cells (**Fig. S4A**). Co-cultivation led to time- and concentration-dependent EV formation (**Fig. S4B**); however, no substantial difference in EV size was observed (**Fig. S4C**). When the Sp-HUS and WT strains were compared at the same apical bacterial concentration, the formation of EVs in the basolateral compartment was higher for the Sp-HUS strain (**Fig. S4D**).

Pneumococcal EVs contain transmembrane proteins and lipoproteins(40), and putative functions have been attributed to Sp-EVs based on their cargo(15). Analysis of the protein composition of Sp-EVs by mass spectrometry analysis revealed that BipA, a 50S ribosomal subunit assembly factor, was the most abundant protein identified in Sp-HUS EVs, followed by DeoD, a purine nucleoside phosphorylase (**Fig. 2F and Table 2**). The fatty acid biosynthesis proteins, FabG and Nox(41), and several proteins related to sugar utilization (e.g., GlgD, GlpK, Gnd, GmpA, and PtsI) (42, 43) were also found at higher abundance in the EVs of the Sp-HUS strain than in those of the WT. Additionally, two choline-binding proteins were more abundant in the Sp-HUS EVs: LytC and the Pneumococcal Surface Protein A (PspA), an immune evasion protein (44–46).

**Table 2.** Proteins with higher abundance in Sp-HUS EVsthan WT EVs.

| Protein ID | Protein name | Gene name and locus |
| --- | --- | --- |
| A0A0H2ZQ39 | 50S ribosomal subunit assembly factor **BipA** (EC 3.6.5.-) (GTP-binding protein BipA) | bipA SPD_0593 |
| Q04L76 | Purine nucleoside phosphorylase DeoD-type (**PNP**) (EC 2.4.2.1) | deoD SPD_0730 |
| A0A0H2ZMC0 | 3-oxoacyl-[acyl-carrier-protein] reductase (EC 1.1.1.100) | fabG SPD_0384 |
| A0A0H2ZNV9 | Glucose-1-phosphate adenyltransferase, **GlgD** subunit (EC 2.7.7.27) | glgD SPD_1007 |
| Q04HZ0 | Glycerol kinase (EC 2.7.1.30) (ATP:glycerol 3-phosphotransferase) (Glycerokinase) (**GK**) | glpK SPD_2013 |
| A0A0H2ZLA5 | 6-phosphogluconate dehydrogenase, decarboxylating (EC 1.1.1.44) | gnd SPD_0343 |
| Q04JB4 | 2,3-bisphosphoglycerate-dependent phosphoglycerate mutase (BPG-dependent PGAM) (PGAM) (Phosphoglyceromutase) (**dPGM**) (EC 5.4.2.11) | gpmA SPD_1468 |
| A0A0H2ZNC0 | 1,4-beta-N-acetylmuramidase, putative (EC 3.2.1.96) | lytC SPD_1403 |
| A0A0H2ZPB4 | NADH oxidase (EC 1.6.99.3) | nox SPD_1298 |
| Q04MI4 | Proline--tRNA ligase (EC 6.1.1.15) (Prolyl-tRNA synthetase) (ProRS) | proS SPD_0246 |
| A0A0H2ZMX6;A0A0H2ZLS0 | Pneumococcal surface protein A | pspA SPD_0126 |
| A0A0H2ZMY1 | Phosphoenolpyruvate-protein phosphotransferase (EC 2.7.3.9) (Phosphotransferase system, enzyme I) | ptsI SPD_1039 |
| A0A0H2ZMK9 | Oxidoreductase, pyridine nucleotide-disulfide, class I | SPD_1415 |
| A0A0H2ZN12 | Putative oxidoreductase (EC 1.-.-.-) | SPD_1590 |
| A0A0H2ZNN3 | General stress protein 24, putative | SPD_1644 |

Proteins found in the Sp-EVs, from both Sp-HUS and WT, are shown in **Table S1**. Choline-binding proteins (CbpC, CbpF, and PcpA), as well as the penicillin-binding protein 1A (Pbp1A) and the pore-forming toxin pneumolysin (Ply), were found in both EV fractions. Several cell division-related proteins, e.g., DivIVA, DnaA, EzrA, FtsA, FtsE, FtsH, FtsX, and FtsZ, and the capsular polysaccharide biosynthesis protein CpsC were also found at similar levels in EVs from both strains. These findings suggest that the biogenesis of pneumococcal EVs is potentially site-specific, occurring preferably at bacterial septa.

### Sp-HUS EVs do not hemolyze red blood cells but strongly bind to human endothelial cells

Human erythrocyte lysis and human endothelial cell viability assays were performed with purified Sp-EVs. Neither Sp-HUS EVs nor WT-EVs lysed erythrocytes (**Fig. 3A**) or affected HUVEC viability (**Fig. 3B**). Next, the direct interaction of Sp-EVs with human endothelial cells was evaluated at the single-cell level by confocal laser-scanning microscopy (CLSM). The expression of the Platelet Endothelial Cell Adhesion Molecule-1 (PECAM-1-FITC, green fluorescence) was assessed in HUVECs challenged with either Sp-HUS EVs or WT-EVs. PECAM-1 is an endothelial cell surface protein known to act as a receptor for pneumococci adhesion to the endothelium, and to be upregulated upon infection and stimulation by pathogen-associated molecules(47–49). Representative microscopy images (**Fig. 3C**) showed that the PECAM-1-FITC signal decreased in the presence of Sp-HUS EVs. The presence of WT-EVs led to heterogeneous PECAM-1-FITC expression, with some HUVECs highly expressing this molecule, whereas others exhibited the baseline FITC level. The semi-quantification of PECAM-1 signal corroborated these findings, in particular the heterogeneous activation profile associated with WT-EVs (**Fig. 3D**). Since Sp-EVs (both WT and HUS EVs) induced lower PECAM-1 expression than the control, it indicates either that Sp-EVs do not activate endothelial cells or that they repress the expression of endothelial adhesion molecules. It was demonstrated that Sp-HUS EVs and WT-EVs negatively affected the expression of the Intercellular Adhesion Molecule-1 (ICAM-1) to a similar extent (**Fig S5**).

**Figure 3.**
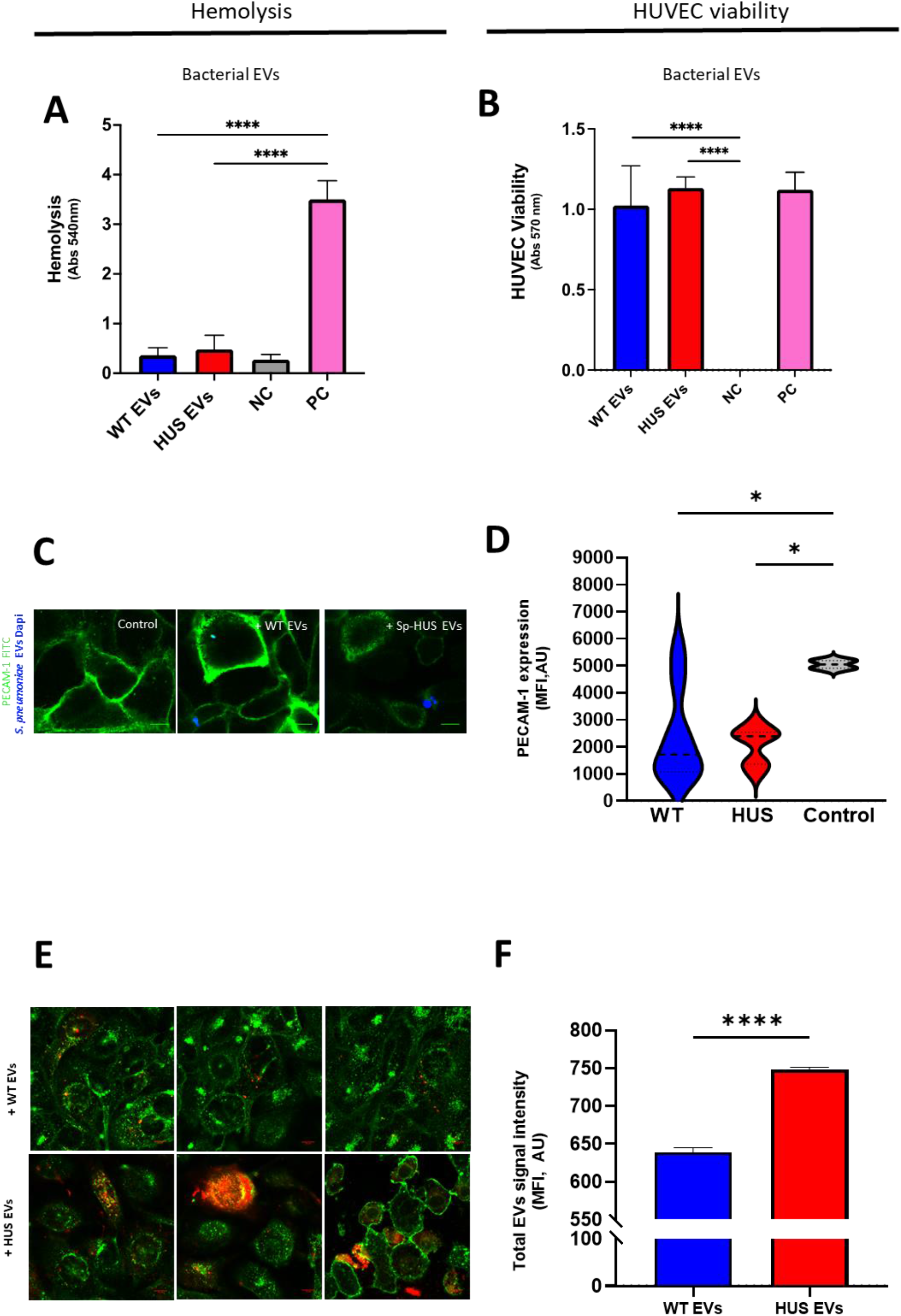
Sp-HUS EVs host-cellular interactions. **(A)** The hemolytic activity of WT and Sp-HUS EVs was tested on human red blood cells, similarly to Fig. 1A. (**B**) HUVECs viability was evaluated after incubation with WT or Sp-HUS EVs by CTB assay, is a similar fashion as Fig. 1C. (**C**) Representative CLSM images of endothelial cells (FITC-labelled anti-PECAM-1 antibody, green) interaction with WT or Sp-HUS EVs (pre-stained with DAPI, blue). HUVECs in growth medium (DMEM) were taken as control. Scale bar = 10 μm. **(D)** Violin plots of the fluorescence intensity of PECAM-1 signal (green) expressed in arbitrary units (AU). **(E)** Representative CLSM images of WT and Sp-HUS EVs (pre-stained with DiD, red) internalization by endothelial cells (WGA CF^®^488A, green). Scale bar = 10 μm. **(F)**Fluorescence intensity of WT (blue bar) and Sp-HUS (red bar) EVs signal expressed in arbitrary units (AU). Means are shown from biological triplicates. Mean ± SD.

Internalization of Sp-EVs by HUVECs was assayed and representative microscopy images highlighted the retention of EV clusters at different locations on HUVEC cells (**Fig. 3E**). Sp-HUS EVs bound to endothelial cells and prominent intracellular colored (red-to-yellow) clusters indicated the internalization of EVs. However, at the resolution level achieved, the precise cytoplasmic localization of these clusters was not possible. The increased (by approximately 14%) intracellular staining of HUVECs by Sp-HUS EVs, compared with that of the WT-EVs, indicated a higher level of Sp-HUS EV internalization (**Fig. 3F**).

In summary, Sp-EVs are non-cytotoxic towards human erythrocytes or endothelial cells, but downregulate endothelial adhesion molecule expression. However, EVs isolated from the HUS strain bound more strongly to endothelial cells than those of the reference strain, identifying this as a relevant feature to consider in the study of HUS infection.

### Sp-HUS EVs elicit cytokine and chemokine release from human monocytes

The capacity of Sp-EVs to elicit an innate immune response in the host was tested by measuring cytokine and chemokine expression quantitative real-time PCR (qRT-PCR; **Fig. 4A**). The transcription of several cytokines and chemokines (IL-1ß, IL-6, TNF-α, CXCL10, Serpin E1, and CCL2) was higher in monocytes incubated with Sp-EVs than in untreated monocytes (**Fig. 4B-G**). Moreover, Sp-HUS EVs induced greater IL-1 ß transcription than did WT-EVs (**Fig. 4B**). For the other tested cytokines, the transcriptional response in monocytes appeared to be essentially induced at the same level by both Sp-HUS EVs and WT-EVs (**Fig. 4C-G)**. Monocyte supernatants were assayed for the presence of cytokines and chemokines by proteome array (**Fig. 4A**). Sp-EVs induced secretion of several cytokines/chemokines from monocytes, including CCL2, CCL3, CXCL1, CXCL10, IL-6, and Serpin E1 (**Fig. 4H**). The proteome array results are in accordance with the transcriptional profile described above for selected cytokines (**Fig.4C-G**), where upregulation was observed after monocyte incubation with Sp-EVs. However, the expression of most of the cytokines/chemokines (CCL2, CCL3, CXCL1, IL-6, and Serpin E1) was higher in response to Sp-HUS EVs than WT-EVs (**Fig. 4H**), suggesting that extracellular vesicles isolated from the HUS strain elicit a stronger innate immune response compared with the reference strain-derived EVs. The production of IL-6 and TNF-α (two major pro-inflammatory molecules) by human monocytes was further investigated at the protein level by an enzyme-linked immunosorbent assay (ELISA), which has a higher sensitivity than the proteome array (**Fig. 4I-J**). The results at the protein level of IL-6 and TNF-α were consistent with their transcriptional levels; their production was induced by both WT and Sp-HUS EVs.

**Figure 4.**
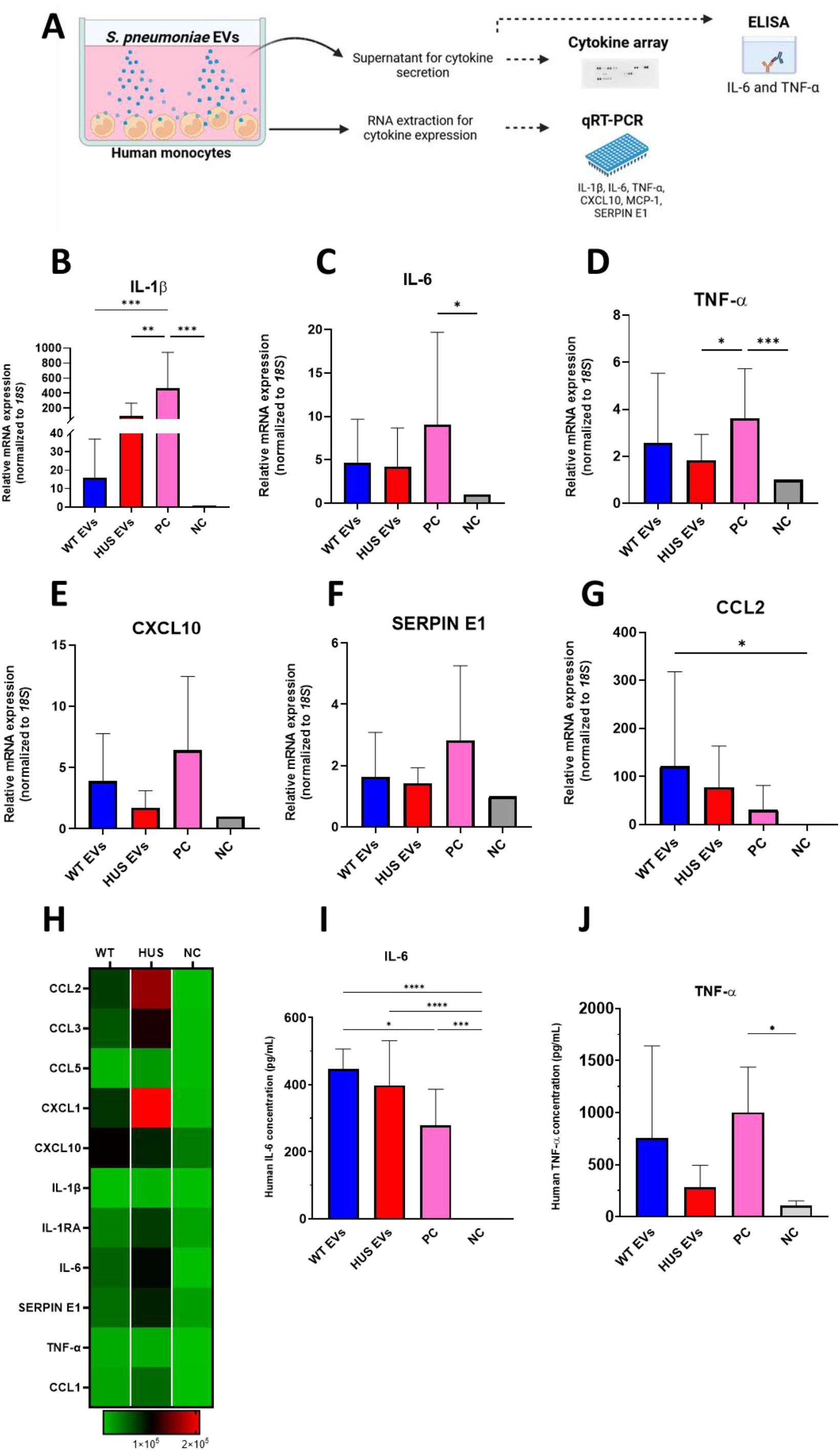
Cytokine expression and production by human monocytes in response to pneumococcal EVs. **(A)** Schematic representation of experiments performed to analyze EVs-induced cytokine expression and production by human monocytes. Monocytes from healthy donors were incubated with WT or Sp-HUS EVs. (**B-G**) Total RNA extracted from monocytes was used to assess transcriptional levels of IL-1β, IL-6, TNF-α, CXCL10, SERPIN E1 and CCl2 by qRT-PCR. **(H)**Supernatants were used to detect cytokine production using a Proteome Profiler Human Cytokine Array. Heat map shows 11 cytokines deregulated in monocytes. **(I, J)**IL-6 and TNF-α production by monocytes quantified by ELISA. In all experiments, untreated monocytes and zymosan treatment (100 μg/mL) were considered the negative (grey bar) and positive (pink bar) controls, respectively. Means are shown from several healthy human donors. Mean ± SD.

Taken together, the results show that EVs isolated from *S. pneumoniae* promote pro-inflammatory cytokine and chemokine transcription and translation in human monocytes, with a more pronounced effect seen with Sp-HUS EVs. The capacity of Sp-HUS EVs for eliciting cytokine production in monocytes, part of the first innate immune response, is an interesting aspect to consider in the understanding of the host immune reaction towards Sp-HUS strain and support the important role that *S. pneumoniae* EVs play in the designing of a vaccine for immunization.

## Discussion

The pathophysiology of the pneumococcal-mediated kidney disease, hemolytic uremic syndrome (Sp-HUS), remains unclear. In this study, we identified pathologic differences between a *S. pneumoniae* HUS strain and a WT strain with regard to their extracellular vesicles (EVs).

Kidney injuries derived from *S. pneumoniae* infections range from proteinuria to acute kidney failure(50, 51). The clinical Sp-HUS strain mediated endothelial damage (**Fig. 1**). Previous reports on Sp-HUS pathogenesis have demonstrated the crucial role played by pneumococcal neuraminidase in exposing the Thomsen-Friedenreich antigen on the surface of the host cell membranes and thus generating damage(52, 53). *NanA* and *nanB*, two neuraminidase genes, were upregulated in the Sp-HUS strain, which aligns with previous knowledge on HUS pathophysiology. Sp-HUS strain might be better adapted to anaerobic growth, including during blood infection where oxygen is less available, as supported by the downregulation of *sufB* and upregulation of *nrdDG*(54–56). However, changes in the transcriptome do not always translate to changes at the proteome level(57), thus conclusions should be carefully drawn, and further studies on the metabolome of HUS clinical strains are necessary to confirm these observations.

Host-derived EVs are associated with STEC-HUS pathology(24). Ståhl *et al.* described a novel mechanism of transfer of a bacterial virulence factor (Stx), attached to blood cell-derived microvesicles, to kidney glomerular endothelial cells(58). This study emphasized the usefulness of microvesicles in mediating the circulation of bacterial toxins, which may lead to immune evasion and ultimately to cellular renal damage. From the results presented in this work, it can be speculated that virulence proteins could be loaded into EVs and act locally, outside the alveoli, in a similar way to that observed for TatD, a pneumococcal endo-deoxyribonuclease involved in neutrophil extracellular trap evasion(59). This potential site-directed and cargo-specific capacity of Sp-HUS EVs would allow damage to the kidney even in the absence of bacteria, as EVs could diffuse into the blood stream and eventually unload their toxic cargo onto kidney endothelial cells.

Sp-HUS EVs exhibited growth-dependent size heterogeneity (**Fig. 2**). Size heterogeneity has been observed for eukaryotic EVs, where it relates to divergent roles in cancer biology(60), and, additionally, EV subpopulations frequently display different cargo(16, 18)(61). We were also intrigued by this heterogeneity and so separated different, small subpopulations by size exclusion chromatography and tested each fraction for its cytotoxicity towards human endothelial cells. Our preliminary data did not show any difference between the fractions (data not shown); however, the separation protocol needs to be optimized to perform a more fine-tuned separation of the small subpopulations. The differential size and concentration observed for Sp-HUS EVs might also relate to their distinct biogenesis.

Proteomic analysis (**Fig. 2F** and **Table 2**) revealed abundant sugar utilization systems proteins in Sp-HUS EVs. These included PtsI, an enzyme belonging to the phosphotransferase system (PTS), and GlgD, a transferase related to glucose metabolism, which were previously shown to be overexpressed in *S. pneumoniae* D39 grown in mannose and mucin, respectively(42, 62). The ability to optimize growth in different sugars is a strategy utilized by several pathogens during host interaction in order to exploit the prevailing environmental conditions(63). The intricate metabolic rearrangements involving glucose and mannose should be a focus for future research. Additionally, the presence of DivIVA, EzrA, and LytC in EVs suggests a possible septal origin for EV formation, as earlier hypothesized by Greenawwalt in the 1970s(64).

Sp-HUS EVs attached to and were internalized by human endothelial cells, in addition to eliciting a pro-inflammatory innate immune response in human monocytes (**Fig. 3** and **Fig. 4**). In previous studies, it was observed that Sp-EVs are promptly internalized by macrophages(16, 18) and epithelial cells(36). In those studies, depending on the pneumococcal strain assayed and on the source of the immune cells, several cytokines had altered expression, including TNF-α, IL-6, IL-10, and IL-1β. Sp-HUS EVs induced higher production of CCL2, CCL3, CXCL1, and CXCL10. Together with IL-6 and Serpin E1, these chemokines are involved in promoting inflammation (65–67). Despite inducing cytokine release, Sp-HUS EVs had a negligible cytotoxic effect on the host cells, corroborating the results of previous studies(11, 68, 69), which strengthens their potential use as immunization tools. Moreover, the presence of PspA and AliA in Sp-EVs (**Table S1**) has previously been associated with high levels of effective protection against *S. pneumoniae*, as shown by reduced bacterial loads in a murine model of pneumococcal colonization(70, 71).

Even though vaccination against this pathogen is available, its efficacy strongly depends on the prevalent pneumococcal serotypes (72, 73). Pneumococcal serotypes are defined by the biochemical structure of their polysaccharide capsule; they are of greatest relevance in the rollout of infection and more than 100 serotypes have been described to date(74). The prominent serotypes of Sp-HUS, before the introduction of the pneumococcal vaccine, were 3, 6B, 8, 9V, 14, 19, and 23F(5). Soon after the introduction of the 7- and 13-valent pneumococcal protein conjugate vaccines in 2000 and 2010, respectively, there was a shift of Sp-HUS-associated serotypes to those that were not covered by the vaccines. Studies carried out in the USA and the UK reported that Sp-HUS cases were mainly caused by serotypes 1, 3, 7F, and, most abundantly, 19A(75, 76). Multiplex PCR(77) revealed that the clinical Sp-HUS strain belongs to the 19A serotype group, precisely the serotype frequently observed after the introduction of protein conjugate vaccination (**Fig. S6**). Initial screens of several Sp-HUS strains (**Fig. 1A** and **B**) showed a high degree of phenotypic heterogeneity, which can be explained, in part, by their potentially different serotypes and, additionally, by unknown specific patient-related issues (age, gender, co-morbidities, medication, etc.).

In conclusion, we showed that substantial differences exist between the Sp-EVs produced by a clinical HUS isolate and those produced by a reference pneumococcal strain. These differences included size and concentration of EVs, their protein cargo, and their ability to evoke inflammatory responses in the host. Sp-HUS EVs might also be carriers of toxins that specifically target the kidney endothelial cells and allow bacteria to avoid direct contact with blood-circulating innate immune cells. Based on this initial characterization, we suggest that Sp-HUS EVs might be good candidates for Sp-HUS diagnosis as they can be easily isolated from the blood or urine of patients and pneumococcal proteins, e.g., BipA and PspA, could act as specific, likely selective Sp-HUS EV markers. Earlier diagnosis of this kidney disease would allow prompter therapeutic intervention, which could prevent the development of more serious outcomes for the patient.

## Material and Methods

### Bacterial strains and growth conditions

The pathogenic strain *Streptococcus pneumoniae* D39 was used as the reference strain(78). Clinical pneumococci strains, isolated from patients with HUS (HUS strains), were obtained from PD Dr. Med. Giuseppina Sparta, Zürich, Switzerland. All *S. pneumoniae* strains were grown in liquid Todd-Hewitt broth (Roth^®^) supplemented with yeast extract (THY) at 37°C in 5% (v/v) CO_2_. Blood agar plates were prepared from blood agar (VWR^®^) with addition of 5% (v/v) defibrinated sheep blood (Thermo Scientific^®^). Growth was monitored by measuring the optical density at 600 nm (OD_600_).

### Cell culture and cell harvesting

Human umbilical vein endothelial cells (HUVECs, CRL-1730) were cultivated in Dulbecco’s modified Eagle’s medium, DMEM (Lonza^®^), supplemented with 10% (v/v) fetal bovine serum (Biochrom^®^), 6 mmol/L l-glutamine (Lonza^®^), and a mixture of penicillin/streptomycin (100U/100 μg/mL, Sigma^®^) at 37°C in the presence of 5% CO_2_. The fully supplemented DMEM medium is referred to as growth medium.

Adherent human cells were washed with pre-warmed Dulbecco’s phosphate-buffered saline (DPBS; Lonza^®^) and harvested by incubation for 10 min at 37°C with PBS containing trypsin/EDTA (Gibco^®^). Cell detachment was stopped by adding 10 mL growth medium. After centrifugation, the pellet was resuspended in 1 mL growth medium and cells were counted using a cell counter CASY (OLS^®^CASY).

### Hemolysis assay

*S. pneumoniae* strains were grown at 37°C in 5% CO_2_ until mid-logarithmic phase was reached. Aliquots of 100 μL red blood cells were incubated in a 96-well plate together with the same volume of bacterial suspensions in THY, bacterial suspensions in PBS, bacterial supernatants, or Sp-EVs. PBS was used as a negative control and bi-distilled water as a positive control. The plates were incubated at 37°C with slight agitation (300 rpm) for 30 min (the positive control was added only 10 min prior to the end of the incubation). The plates were immediately centrifuged (400 g, 15 min, RT) before the resulting supernatants were transferred to a fresh 96-well plate and their OD at 540 nm was measured.

### Hydrogen peroxide measurement

*S. pneumoniae* strains were grown as previously described, and after centrifugation, supernatants were filtered and immediately assayed for the presence of hydrogen peroxide using the Hydrogen Peroxide Colorimetric Assay (Biocat), following manufacturer instructions. Uninoculated growth medium (THY) was used as the negative control and its value was subtracted from all the absorbances measured.

### Super Resolution-Structured Illumination Microscopy (SR-SIM)

SR-SIM was performed to visualize cell wall staining in single bacterial cells. Briefly, bacteria were grown in THY at 37°C in 5% CO_2_ until mid-exponential phase. After washing with DPBS, bacteria were stained with CF^®^488-conjugated Wheat Germ Agglutinin (WGA; biotium^®^) for 1 h in the same incubation conditions. Bacterial cells were then washed and fixed with 4% (v/v) paraformaldehyde (PFA) solution for 20 min at 4°C. For the SR-SIM imaging, 10 μl of the sample was spotted on 1% agarose pads. The agarose pads were covered with No. 1.5H coverslips (Roth) and stored at 4°C for further imaging. The SR-SIM data were acquired on an Elyra 7 system (Zeiss) equipped with a 63 ×/1.4 NA Plan-Apochromat oil-immersion DIC M27 objective lens (Zeiss), a Piezo stage, and a PCO edge sCMOS camera with 82% QE and a liquid cooling system with 16-bit dynamic range. Using Lattice SIM mode, images were acquired with 13 phases. WGA CF^®^488 was detected with a 488 nm laser and a BP 495-590 emission filter. Super resolution images were computationally reconstructed from the raw data sets using default settings on ZenBlack software (Zeiss). Images were analyzed using the Fiji ImageJ software(79).

### Endothelial cell viability assay

The cytotoxicity of *S. pneumoniae* cells, supernatants, and Sp-EVs towards human endothelial cells was accessed by the CellTiter-Blue^®^ (CTB) Cell Viability Assay (Promega), according to manufacturer’s instructions. HUVECs were seeded in 96-well plates (Thermo Scientific^®^) at a density of 1.5×10^4^ cells/well. Cells were cultivated at 37°C in 5% CO_2_ until confluence was reached. Mid-exponentially-grown *S. pneumoniae* strains, Sp-supernatants, or Sp-EVs were incubated with HUVEC for 1 h under the same growth conditions. Tert-butyl hydroperoxide (400 μM) and DMEM were used as negative and positive cell viability controls, respectively. Subsequently, gentamicin (500 mg/mL) was added to the medium for 1 h to kill any extracellular bacteria that would otherwise contribute to the cell viability measurements. HUVECs were then washed with pre-warmed DPBS and CTB (100 μL) was added to each well. Following incubation for 16 h at 37 °C in 5% CO_2_, the absorbance of each well was measured at 570 nm using a Tecan^®^ Safire 2 microplate reader. In this assay, metabolically viable endothelial cells convert the redox dye (resazurin) into a fluorescent end product (resorufin). Statistical analysis was performed using Prism version 9 for Windows (GraphPad Software, La Jolla, CA).

### RNA seq

Total RNA was isolated from bacterial cells using a universal RNA purification kit (roboklon) and a subsequent clean up step was performed using a Cleanup kit (Monarch^®^RNA), both according to the manufacturer’s instructions. cDNA libraries were prepared by vertis Biotechnology AG (Freising, Germany). The ribodepleted RNA samples were first fragmented using ultrasound (1 pulse of 30 s at 4°C), and then an oligonucleotide adapter was ligated to the 3’ end of the RNA molecules. First-strand cDNA synthesis was performed using M-MLV reverse transcriptase and the 3 ‘ adapter as primer. The first-strand cDNA was purified and the 5’ Illumina TruSeq sequencing adapter was ligated to the 3’ end of the antisense cDNA. The resulting cDNA was PCR-amplified to about 10–20 ng/μl using a high fidelity DNA polymerase. The cDNA was purified using the Agencourt AMPure XP kit (Beckman Coulter Genomics) and was analyzed by capillary electrophoresis. The cDNA pool was sequenced on an Illumina NextSeq 500 system using 75 bp read length. The read files in FASTQ format were imported into CLC Genomics Workbench v11 (Qiagen) and trimmed for quality. Reads were mapped to the *S. pneumoniae* reference genome (NCBI accession numbers: NC_008533.2) using the ”bowtie2” tool with standard parameters and quantified using “featureCounts”. Reads counts were normalized using the median of ratios method from DESeq2 package from R(35). Pearson correlation and Principal Component Analysis were performed to ascertain the degree of correlation between replicates. Even though replicates #1 and #2 differed from the other three replicates, differential gene expression analysis could still be conducted. Genes with a |fold change| > 1.5 and an adjusted *p*-value < 0.05 were considered as differentially expressed in the two strains.

### Extracellular vesicle isolation

For bacterial EVs (Sp-EVs), *S. pneumoniae* strains were grown on a solid blood agar plate overnight and single colonies were inoculated into full supplemented DMEM media and grown at 37°C in 5% CO_2_. At mid-logarithmic growth phase, 10 mL aliquots of bacterial culture were taken and centrifuged (4,000 g, 15 min, 4°C). The pellet was discarded and the supernatant was filtered through a 0.45 μm pore membrane (Sartorius) twice in order to obtain cell-free supernatant. The resulting cell-free media was then centrifuged at 100,000 g for 2 h at 4°C using the type 70.1 Ti rotor from Beckam Colter^®^. The resulting vesicle pellets were resuspended in 10 mL sterile PBS and the centrifugation step was repeated for 1 h. Final washed vesicle pellets were resuspended in PBS and stored at −20°C for further analysis. Additionally, EVs were precipitated using ExoQuick-TC (System Biosciences) according to the manufacturer’s protocol. For subsequent use, EVs were slowly thawed on ice.

### Scanning Electron Microscopy (SEM)

For SEM, bacteria were grown as described above, and seeded on 12 mm Ø coverslips (Roth^®^). Cells were fixed for 1 h in 2.5% (v/v) glutaraldehyde in sodium cacodylate buffer (0.1 M, pH 7.0) and washed three times with sodium cacodylate buffer for 20 minutes each. Samples were dehydrated in increasing ethanol concentrations followed by critical point drying using a Leica EM CPD300 Automated Critical Point Dryer (Leica) and finally coated with gold (25 nm) in a Safematic CCU-010 HV Sputter Coating System (Safematic). SEM images were acquired at different magnifications in a Zeiss-LEO 1530 Gemini field-emission scanning electron microscope (Carl Zeiss) at 6–8 kV acceleration voltage and a working distance of 5–7 mm using an InLense secondary electron detector for secondary electron imaging.

### Extracellular vesicles counting

Isolated vesicles were counted using a NS300 dynamic light-scattering microscope (Malvern) fitted with NanoSight NTA 3.2 software. Isolated vesicles were dispersed in 1 mL DPBS and injected though the microscope at 100 (AU) pump flow rate. Videos were captured at 24 fps for three periods of 60 seconds for each sample and analyzed using NanoSight NTA 3.2.

### Confocal laser-scanning microscopy of EV and endothelial cell interactions

For characterization studies, HUVECs and Sp-EVs were co-incubated. After isolating Sp-EVs as described above, vesicles were incubated for 30 minutes at room temperature with 4’,6-diamidino-2-phenylindole, dihydrochloride (DAPI, Biotium^®^) or DiD (Vibrant^TM^ Cell-Labeling solutions, Molecular Probes), washed with PBS, and resuspended in growth medium.

HUVECs were seeded on 18 mm Ø coverslips in the wells of 24-well plates (7 × 10^4^ cells/well), and when confluency was reached, cells were washed with DPBS and processed for either immunoblotting staining or were directly stained with WGA CF^®^488A (Biotium^®^) for 30 minutes at 37°C in 5% CO_2_. For immunoblot staining, HUVECs were first incubated with pre-stained vesicles, blocked (PBS with 0.5% [v/v] Tween20, 0.5% [w/v] Bovine Serum Albumin and 4% [w/v] milk), and further incubated for 1 h at room temperature with FITC-labeled anti-PECAM-1 antibody (Platelet Endothelial Cell Adhesion Molecule-1, Abcam) diluted 1:1000 in blocking buffer before being washed twice with PBS-T. Endothelial cells were incubated with pre-stained vesicles for 2 h (adhesion) or 4 h (internalization) at 37°C in 5% CO_2_, and then fixed with 4% (v/v) paraformaldehyde (PFA) for 20 minutes at 4°C. The imaging was carried out on a Confocal Laser Scanning Microscope (LSM 710, Zeiss^®^) equipped with ZEN 2011 software.

### Sample preparation for proteome analysis

Proteins from EVs were extracted with 1% (w/v) SDS in PBS (5 min, 95°C), reduced with 50 mM DTT (5 min, 95 °C), and diluted with 8 M Urea in 100 mM Tris/HCl, pH 8.0. Buffer exchange and protein digestion was carried out according to the FASP protocol(80). In brief, the reduced proteins were transferred to a 30 kDa Microcon filter unit (YM-30 filter units, Millipore) and centrifuged at 14,000 × g for 20 min in all consecutive steps, and the flow-through was discarded. For washing, 200 μL urea buffer (8 M Urea, 100 mM Tris HCL, pH 8.0) was added and the centrifugation was repeated. Aliquots (100 μL) of alkylation solution (0.1 M iodoacetamide in urea buffer) were added, and samples were incubated for 20 min in the dark and subsequently washed (200 μL 8 M urea buffer) after each of two additional centrifugation steps. Afterwards, samples were centrifuged twice and washed with 200 μL 50 mM ammonium bicarbonate buffer, and proteins were digested by the addition of 0.5 μg trypsin in 50 μL 50 mM ammonium bicarbonate. Proteolytic cleavage was allowed to proceed for 16 h at 37°C and peptides were eluted by centrifugation. Eluted peptides were dried in a SpeedVac (Thermo Fisher) and reconstituted by adding 20 μL of 0.1% (v/v) formic acid in water.

### Mass spectrometry and statistical analysis

Tryptic peptides were analyzed with a Dionex UHPLC (Thermo Scientific) coupled to an Orbitrap Fusion ETD (Thermo Scientific). Peptides were loaded onto a trap column (Acclaim PepMap C18) and separated using a 50 cm analytical column (PepMap RSLC, C18). Full MS1 scans were acquired in the Orbitrap (m/z range 370–1570, quadrupole isolation) at a resolution of 120,000 (full width at half maximum) during a 2.5 h, non-linear gradient ranging from 2% to 90% (v/v) acetonitrile/0.1% (v/v) formic acid. Peptides were fragmented by higher energy collisional dissociation (HCD, 32% collision energy) and max. 20 fragment ion spectra were acquired per cycle in the ion trap in rapid mode (quadrupole isolation m/z window = 1.6). The following conditions were used: spray voltage of 2.0 kV, heated capillary temperature of 275 °C, S-lens RF level = 60%, a maximum automatic gain control (AGC) value of 4×10^5^ counts for MS1 with a maximum ion injection time of 50 ms and a maximum AGC value of 2×10^3^ for MS2, with a maximum ion accumulation time of 35 ms. A dynamic mass exclusion time window of 60 s was set with a 10 ppm maximum mass window. All raw files were searched against reference proteomes of *S. pneumoniae* D39 (version 03.2021) and *Bos taurus* (version 10.2021) with MaxQuant version 1.6.17.0 (Max Planck Institute of Biochemistry, Germany)(81). The default parameters were used or set as follows: enzyme: trypsin, max. 2 missed cleavages; static modification: carbamidomethylation of cysteine residues; variable modifications: methionine oxidation; min. peptide length: 6; max. peptide mass: 7600 Da. Normalization was done in MaxQuant using Label-Free Quantification (LFQ) min. ratio count = 2 (unique and razor peptides) and matching between runs was enabled. PSM (peptide specific matches) and protein FDR was set to 0.01. LFQ values of all samples were loaded into Perseus (version 1.6.2.2)(82). The resulting matrix was reduced as proteins identified as “only identified per site”, ”reverse identified”, and “potential contamination” were discarded.

### Monocyte isolation and co-incubation with EVs

Human monocytes were isolated from the buffy coat of healthy male donors (University Hospital Jena, German Red Cross). The cells were isolated by density gradient centrifugation. Shortly, 30 mL buffy coat was diluted with 5 mL DPBS (BioWhittaker^®^) and carefully poured over 15 mL of Histopaque^®^ (Merck). After centrifugation (500 g, 20 min, RT, no acceleration break), the peripheral blood mononuclear cell (PBMC) ring was collected. Cells were washed with DPBS (100 g, 5 min, RT) and a second density gradient centrifugation step was performed to exclude remaining red blood cells and neutrophils. Finally, cells were poured over 46% (v/v) Percoll^®^ (Merck) diluted with Iscove’s Modified Dulbecco’s Medium (IMDM; Thermo Fisher) to remove lymphocytes and re-centrifuged (500 g, 20 min, RT). The monocytes were collected, washed, resuspended in 1 mL IMDM medium supplemented with 10% (v/v) fetal calf serum (FCS) (termed conditioned medium (CM)), and counted using the cell counter CASY (OLS^®^CASY). Subsequently, 1×10^8^ cells were seeded into a cell culture flask in 15 mL CM and incubated at 37 °C in 5% CO_2_ for 2 h to allow monocyte adherence to the plastic surface. Afterwards, the cell layer was washed three times with DPBS to remove all non-adherent cells and the monocytes were incubated overnight at 37 °C in 5% CO_2_.

EVs isolated as described above were directly added to the monocyte layer and incubated for 24 h at 37 °C in 5%CO_2_ to assess cytokine gene transcription and translation in response to Sp-EVs. Controls for the assays included zymosan-treated monocytes (100 μg/mL) as the positive control and monocytes in growth medium as the negative control.

### Quantitative Real-Time PCR (qRT-PCR) of cytokine genes

Monocytes were challenged with Sp-EVs as described above, after which their total RNA was extracted, using the Universal RNA Purification Kit (EURx^®^ Roboklon) according to the manufacturer’s instructions, and quantified using an ND-1000 spectrophotometer (Nanodrop).

The expression of genes encoding 18S ribosomal RNA (rRNA), IL-6, IL-1ß, TNF-α, monocyte chemoattractant protein (MCP)-1, C-X-C motif chemokine ligand 10 (CXCL10), and SERPIN E1 was assessed by qRT-PCR. cDNA was synthesized from 1 μg total RNA from each stimulated sample using High-Capacity cDNA Reverse Transcription Kit (Applied Biosystems) according to the manufacturer’s instructions. Gene expression was quantified on QuantStudio 6 PRO Real-Time PCR System using PowerUp™ SYBR™ Green Master Mix (Applied Biosystems) and gene-specific primers (**Table S4**). Relative gene expression levels were normalized against the 18S rRNA housekeeping gene expression and calculated using the 2-ΔΔCt method.

### Detection of monocyte cytokine production in response to Sp-EVs

After incubating monocytes with Sp-EVs as described before, cell-free supernatant was collected and a semiquantitative immunosorbent assessment of cytokine expression was performed using a Proteome Profiler Human Cytokine Array Kit (R&D Systems) according to the manufacturer’s instructions. Briefly, after blocking, membranes were incubated with the samples and detection antibody cocktail overnight at 4°C. The membranes were then washed and incubated with streptavidin-horseradish peroxidase (HRP) at room temperature for 30 minutes, and signals were detected using chemiluminescent detection reagents (CheLuminate-HRP PicoDetect, PanReac AppliChem). The pixel density of each spot was measured using the image analysis software Fiji(79). The level of interleukin (IL)-6 and tumor necrosis factor (TNF)-α was also quantified by Human IL-6 and TNF-α ELISA Kits (ImmunoTools), respectively.

### Statistical analysis

Unless otherwise stated, statistical significance was determined by ordinary one-way analysis of variance (ANOVA) test with a Bonferroni multiple comparisons test. Probability values (*p*-values) were defined as follows: ns, *, *p* ≤ 0.05; **, *p* ≤ 0.01; ***, *p* ≤ 0.001; ****, *p* ≤ 0.0001.

## Acknowledgements

The work of the authors is supported by the Collaborative Research Center, FungiNet (project C6 (PFZ)) Deutsche Forschungsgemeinschaft (DFG). SH received funding by the Deutsche Forschungsgemeinschaft DFG HA 3125/5-2 and DFG-GRK 2719.

We thank the Microverse Imaging Center for providing microscope facility support for data acquisition. The ELYRA 7 was funded by the Free State of Thuringia with grant number 2019 FGI 0003. The Microverse Imaging Center is funded by the Deutsche Forschungsgemeinschaft (DFG, German Research Foundation) under Germany’s Excellence Strategy - EXC 2051 - Project-ID 390713860.

## Author Contributions

MB and CV conceived and designed the experiments. MB, LT, LZ, SL and MMM performed the experiments. MB, CV, BH, YB, LT, LZ and MMM analyzed the data. MW, AJ, HS, SH, MTF and PZ contributed with reagents, materials, and analysis tools. CV, MB, BH, YB, MMM and PZ wrote the manuscript. CV, MW, AJ, HS, SH, MTF and PZ corrected the manuscript.

